# The tRNA thiolation-mediated translational control is essential for plant immunity

**DOI:** 10.1101/2022.02.13.480182

**Authors:** Xueao Zheng, Hanchen Chen, Zhiping Deng, Yujing Wu, Linlin Zhong, Chong Wu, Qiansi Chen, Shunping Yan

**Affiliations:** Hubei Hongshan Laboratory, Wuhan, 430070, China; College of Life Science and Technology, Huazhong Agricultural University, Wuhan, Hubei 430070, China; Shenzhen Institute of Nutrition and Health, Huazhong Agricultural University, Shenzhen 518000, China; Shenzhen Branch, Guangdong Laboratory for Lingnan Modern Agriculture, Shenzhen 518000, China; Agricultural Genomics Institute at Shenzhen, Chinese Academy of Agricultural Sciences, Shenzhen 518000, China; Zhengzhou Tobacco Research Institute of CNTC, No.2 Fengyang Street, Zhengzhou, Henan Province, 450001, China; State Key Laboratory for Managing Biotic and Chemical Threats to the Quality and Safety of Agro-products, Institute of Virology and Biotechnology, Zhejiang Academy of Agricultural Sciences, Hangzhou 310021, China; Key Laboratory of Horticultural Plant Biology, Ministry of Education, College of Horticulture and Forestry Sciences, Huazhong Agricultural University, Wuhan 430070, China

**Keywords:** plant immunity, translation, tRNA thiolation, NPR1, *Arabidopsis*

## Abstract

Plants have evolved sophisticated mechanisms to regulate gene expression to activate immune responses against pathogen infections. However, how the translation system contributes to plant immunity is largely unknown. The evolutionarily conserved thiolation modification of tRNA ensures efficient decoding during translation. Here we show that tRNA thiolation is required for plant immunity in *Arabidopsis*. We identify a *cgb Arabidopsis* mutant, which is hyper-susceptible to the pathogen *Pseudomonas syringae. CGB* encodes ROL5, a homolog of yeast NCS6 required for tRNA thiolation. ROL5 physically interacts with CTU2, a homolog of yeast NCS2. Mutations in either *ROL5* or *CTU2* result in loss of tRNA thiolation. Further analyses reveal that both transcriptional reprogramming and translational reprogramming during immune responses are compromised in *cgb.* Notably, the translation of the salicylic acid receptor NPR1 is reduced in *cgb*, resulting in reduced salicylic acid signaling. Our study not only reveals a new regulatory mechanism for plant immunity but also uncovers a new biological function of tRNA thiolation.

## Introduction

As a sessile organism, plants are frequently infected by different pathogens, which greatly affect plant growth and development, and cause a tremendous loss in agriculture (Jones and Dangl, 2006; Yan et al., 2013; Spoel and Dong, 2012). To defend against pathogens, plants have evolved sophisticated immune mechanisms. One of the essential immune regulators is the phytohormone salicylic acid (SA), which is considered an immune hormone and plays a central role in immune responses (Zhou and Zhang, 2020; Corina Vlot et al., 2009; Yan and Dong, 2014; Peng et al., 2021). Upon pathogen infection, the biosynthesis of SA is dramatically induced. Plants defective in SA biosynthesis or SA signaling are hyper-susceptible to pathogens (Rekhter et al., 2019; Cao et al., 1997). Recent studies have shown that NONEXPRESSER OF PR GENES 1 (NPR1) and its homologs NPR3 and NPR4 are SA receptors (Fu et al., 2012; Wu et al., 2012; Ding et al., 2018). While NPR1 is a positive regulator of SA signaling, NPR3 and NPR4 are negative regulators.

Immune responses involve massive changes in gene expression at transcription, post-transcription, translation, and post-translation levels. Compared with other regulatory mechanisms, the translation regulation mechanism is less well-studied. However, accumulating evidence suggested the importance of translation regulation in plant immune responses. It is reported that both the pattern-triggered immunity (PTI) and effector-triggered immunity (ETI) involves translational reprogramming (Xu et al., 2017; Yoo et al., 2019). And PABP/purine-rich motif is an initiation module for PTI-associated translation (Wang et al., 2022) and CDC123, an ATP-grasp protein, is a key activator of ETI-associated translation (Chen et al., 2023).

During translation, the code information of mRNA is decoded by transfer RNA (tRNA) molecules, which carry different amino acids. In this sense, the tRNA molecules function as deliverers of the building blocks for translation. The decoding efficiency of tRNA is affected by tRNA abundance, tRNA modification, aminoacyl-tRNA synthetase, amino acid abundance, and elongation factors, among which tRNA modification is emerging as a key regulator during elongation (Torres et al., 2014; Delaunay et al., 2016; Schaffrath and Leidel, 2017).

Currently, more than 150 different tRNA modifications have been identified (Agris et al., 2018). Among them, the 5-methoxycarbonylmethyl-2-thiouridine of uridine at wobble nucleotide (mcm^5^s^2^U) is highly conserved in all eukaryotes. The mcm^5^s^2^U modification is present in the wobble position of tRNA-Lys(UUU), tRNA-Gln(UUG), and tRNA-Glu(UUC) (Huang et al., 2005; Lu et al., 2005; Sen and Ghosh, 1976). In budding yeast (*Saccharomyces cerevisiae*), the 5-methoxycarbonylmethyl of uridine (mcm^5^U) is catalyzed by the Elongator Protein Complex (ELP) and Trm9/112 complex, and the thiolation (s^2^U) is mediated by the URM1 pathway involving URM1, UBA4, NCS2, and NCS6 (Nakai et al., 2004; Noma et al., 2009; Zabel et al., 2008; Leidel et al., 2009). Loss of the mcm^5^s^2^U modification causes ribosome pausing at AAA and CAA codons (Nedialkova and Leidel, 2015; Ranjan and Rodnina, 2017; Rezgui et al., 2013), which results in defective co-translational folding of nascent peptides and protein aggregation, thereby disrupting proteome homeostasis (Rezgui et al., 2013; Nedialkova and Leidel, 2015; Ranjan and Rodnina, 2017). In yeasts, the mcm^5^s^2^U modification was reported to regulate cell cycle, DNA damage repair, and abiotic stress responses (Dewez et al., 2008; Jablonowski et al., 2006; Klassen et al., 2017; Zinshteyn and Gilbert, 2013; Leidel et al., 2009; Nedialkova and Leidel, 2015). In humans, loss of the mcm^5^s^2^U modification causes numerous disorders including severe developmental defects, neurological diseases, tumorigenesis, and cancer metastasis (Shaheen et al., 2019; Pan, 2018; Simpson et al., 2009; Waszak et al., 2020; Torres et al., 2014). In plants, loss of the mcm^5^s^2^U modification was associated with developmental defects and hypersensitivity to heat stress (Nakai et al., 2019; Xu et al., 2020; Leiber et al., 2010). However, it remains unknown whether the mcm^5^s^2^U modification is involved in plant immune responses.

In this study, we found that the mcm^5^s^2^U modification is required for plant immunity. Transcriptome and proteome analyses revealed that the mcm^5^s^2^U modification is essential for the optimal expression of immune-related genes. Especially, we found that SA biosynthesis and signaling is compromised in the mcm^5^s^2^U mutant. Our study not only expands the biological function of tRNA thiolation but also highlights the importance of translation control in plant immunity.

## Results

### ROL5 is required for plant immunity

In a study to test disease resistance of transgenic *Arabidopsis*, we found that one transgenic line was hyper-susceptible to bacterial pathogen *Pseudomonas syringae* pv. *Maculicola* (*Psm*) ES4326. The disease symptom resembled that of *npr1*, in which the master immune regulator NPR1 was mutated (Figure 1A and B). We named this line *cgb* (for Chao Gan Bing; “hyper-susceptible to pathogens” in Chinese). The disease susceptibility of *cgb* was likely due to T-DNA insertion rather than the overexpression of transgene because only one line showed such phenotype. To identify the causal gene of *cgb*, we sequenced its genome using the next-generation sequencing technology, which revealed that there was a T-DNA insertion in the third exon of *ROL5* (AT2G44270; Figure 1C). The insertion was confirmed through genotyping analysis (Figure 1D). In the *cgb* mutant, the transcript of *ROL5* was undetectable, indicating that it was a knock-out mutant (Figure 1E). To confirm that *ROL5* was the *CGB* gene, we carried out a complementation experiment by transforming *ROL5* into the *cgb* mutant. As shown in Figure 1A and 1B, the disease phenotype of the complementation line (*COM*) was similar to that of wild-type (WT). To further confirm this, we generated another allele of *ROL5* mutant, *rol5-c*, using the CRISPR-Cas9 gene-editing approach (Wang et al., 2015). In *rol5-c*, there is a 2-bp deletion in the first exon of *ROL5*, which caused frameshift (Figure 1C). As expected, the *rol5-c* mutant was as susceptible to *Psm* as *cgb* (Figure 1A and 1B). These data strongly suggested that ROL5 is required for plant immunity.

**Figure 1.**
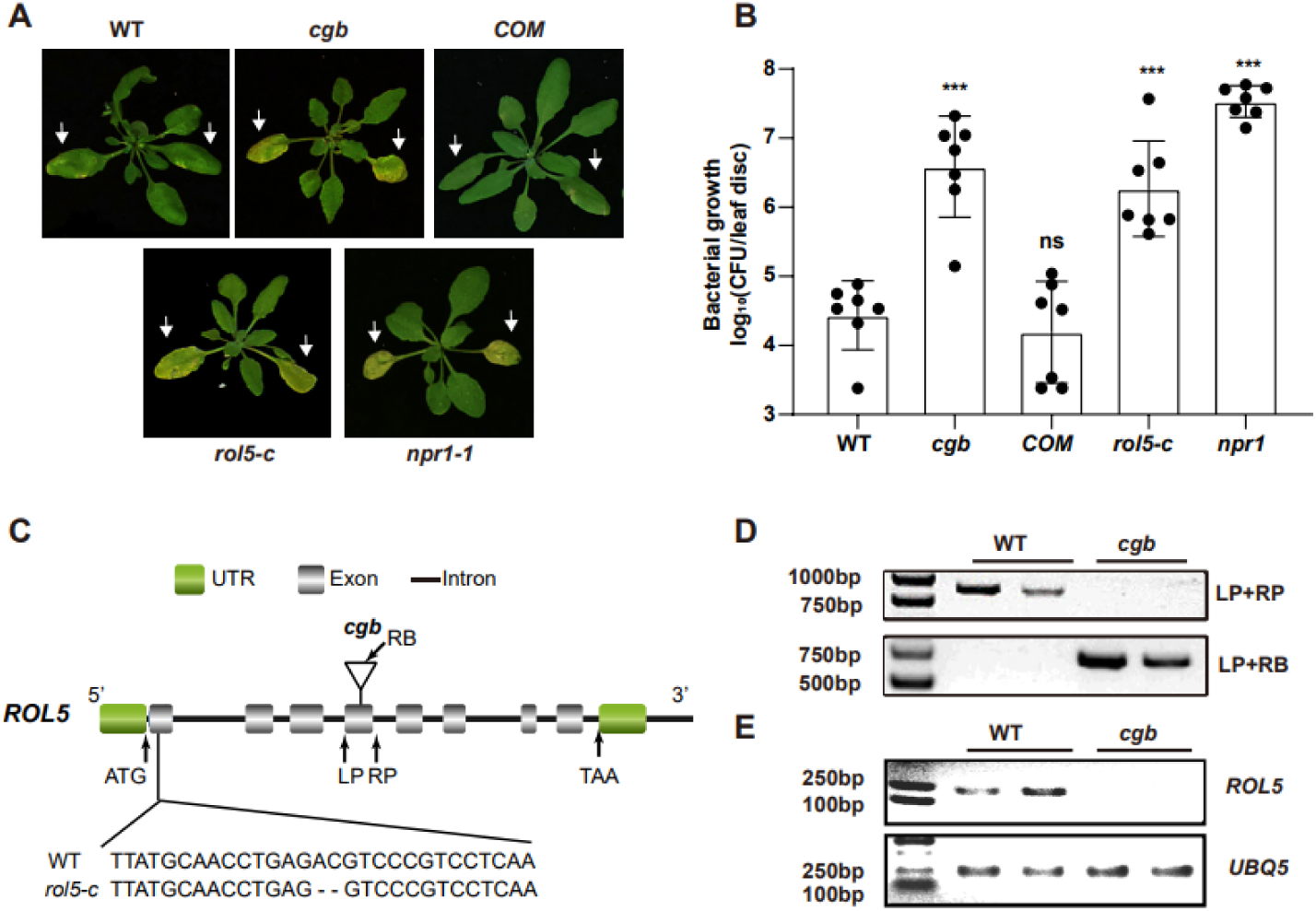
The *rol5* mutants are more susceptible to the bacterial pathogen *Psm* ES4326 than wild-type (WT). (A) The photo of *Arabidopsis* 3 days after infection. The arrows indicate the leaves inoculated with *Psm* ES4326 (OD_600_ = 0.0002). *cgb* and *rol5-c* are mutants defective in ROL5. *COM*, the complementation line of *cgb*. *npr1-1* serves as a positive control. (B) The growth of *Psm* ES4326. CFU, colony-forming unit. Error bars represent 95% confidence intervals (n = 7). Statistical significance was determined by (Student’s t-test, two-tailed). ***, p < 0.001; ns, not significant. (C) A schematic diagram showing the site of T-DNA insertion in *cgb* and the deleted nucleotides in *rol5-c*.(D) The genotyping results using the primers indicated in C. (E) The transcript of *ROL5* is not detectable in *cgb*. *UBQ5* serves as an internal reference gene.

### ROL5 interacts with CTU2 in *Arabidopsis*

ROL5 is a homolog of yeast NCS6 (Leiber et al., 2010), which forms a protein complex with NCS2 to catalyze mcm^5^s^2^U34 (Figure 2A). The NCS2 homolog in *Arabidopsis* is CTU2 (Philipp et al., 2014). To test whether ROL5 interacts with CTU2, we first performed yeast-two-hybrid assays. As shown in Figure 2B, only when ROL5 and CTU2 were co-expressed, the yeasts could grow on the selective medium, indicating that ROL5 interacts with CTU2 in yeast. To test whether they can interact in *vivo*, we carried out split luciferase assays in *Nicotiana benthamiana*. ROL5 was fused with the N-terminal half of luciferase (nLUC) and CTU2 was fused with the C-terminal half of luciferase (cLUC). An interaction between two proteins brings the two halves of luciferase together, leading to enzymatic activity and production of luminescence that is detectable using a hypersensitive CCD camera. As shown in Figure 2C, the luminescence signal could be detected only when ROL5-nLUC and cLUC-CTU2 were co-expressed. We also performed co-immunoprecipitation (CoIP) assays in *N. benthamiana*. When ROL5-FLAG was co-expressed with CTU2-GFP, ROL5-FLAG could be immunoprecipitated by the GFP-Trap beads (Figure 2D). To test whether the interaction is direct, we conducted pull-down assays. GST-CTU2 and ROL5-His proteins were expressed in *Escherichia coli* and were purified using affinity resins. As shown in Figure 2E, ROL5-His could be specifically pulled down by GST-CTU2, but not the GST control, suggesting that ROL5 directly interacts with CTU2.

**Figure 2.**
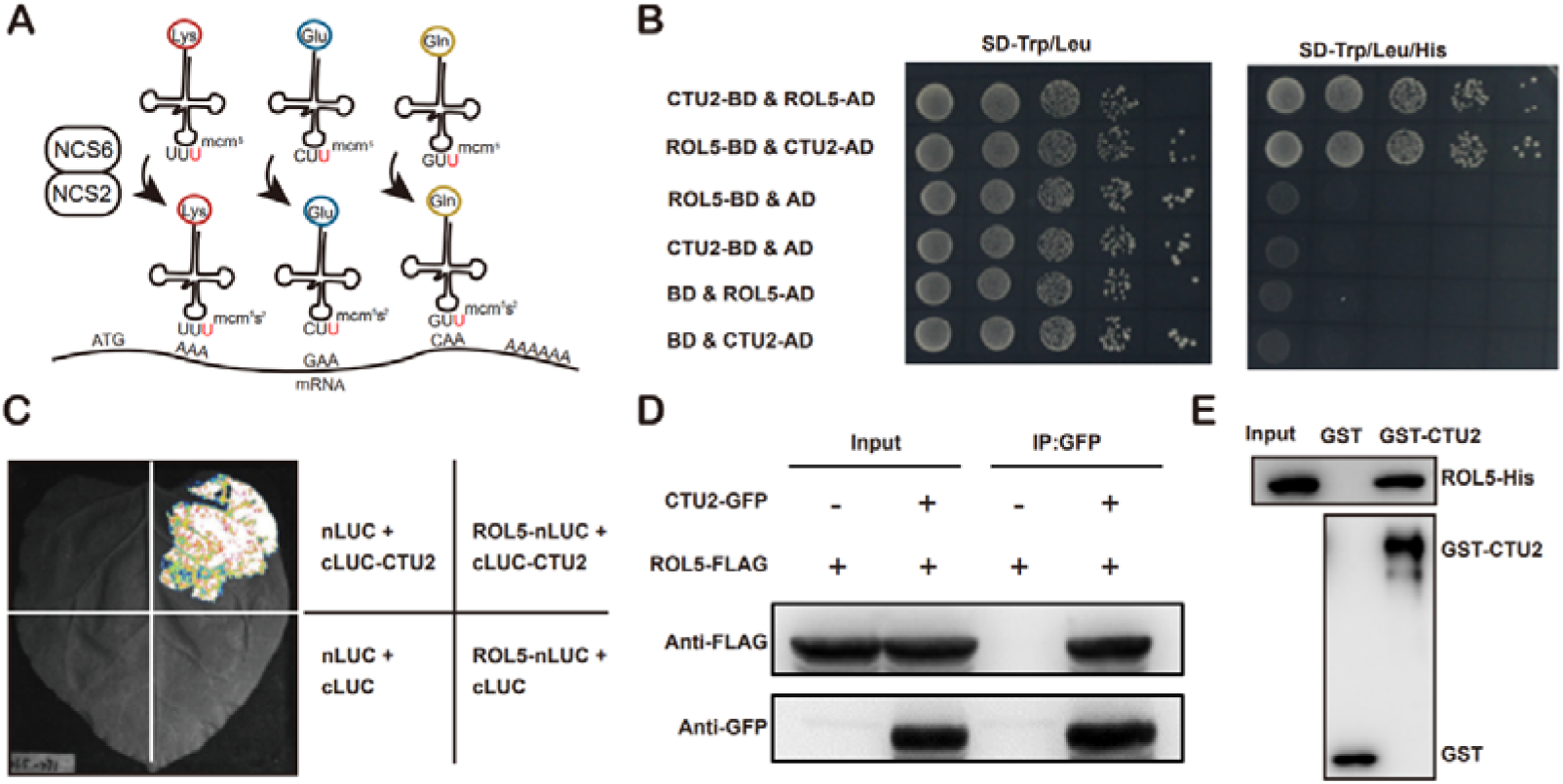
ROL5 interacts with CTU2. (A) A schematic diagram showing the function of ROL5 and CTU2. The ROL5 homolog NCS6 and the CTU2 homolog NCS2 form a complex to catalyze the mcm^5^s^2^U modification at wobble nucleotide of tRNA-Lys (UUU), tRNA-Gln (UUC), and tRNA-Glu (UUG), which pair with the AAA, GAA, and CAA codons in mRNA, respectively. (B) Yeast two-hybrid assay. The yeast growth on the SD-Trp/Leu/His medium indicates interaction. BD, binding domain. AD, activation domain. (C) Split luciferase assay. The proteins were fused to either the C- or N-terminal half of luciferase (cLUC or nLUC) and were transiently expressed in *N. benthamiana*. The luminesce detected by a CCD camera indicates interaction. (D) CoIP assays. CTU2-GFP and/or ROL5-FLAG fusion proteins were expressed in *N. benthamiana*. The protein samples were precipitated by GFP-Trap, followed by western blotting using anti-GFP or anti-FLAG antibodies. (E) The Pull-down assay. The recombinant GST or GST-CTU2 proteins coupled with glutathione beads were used to pull down His-ROL5, followed by western blotting using anti-His or anti-GST antibodies.

### The tRNA thiolation is required for plant immunity

Given that CTU2 interacts with ROL5, we reasoned that the *ctu2* mutant should have similar phenotypes to *rol5* in response to pathogens. To test this, we infected the T-DNA insertion mutant *ctu2-1* with *Psm* ES4326. As expected, the *ctu2-1* mutant is hyper-susceptible to pathogens (Figure 3A and 3B).

**Figure 3.**
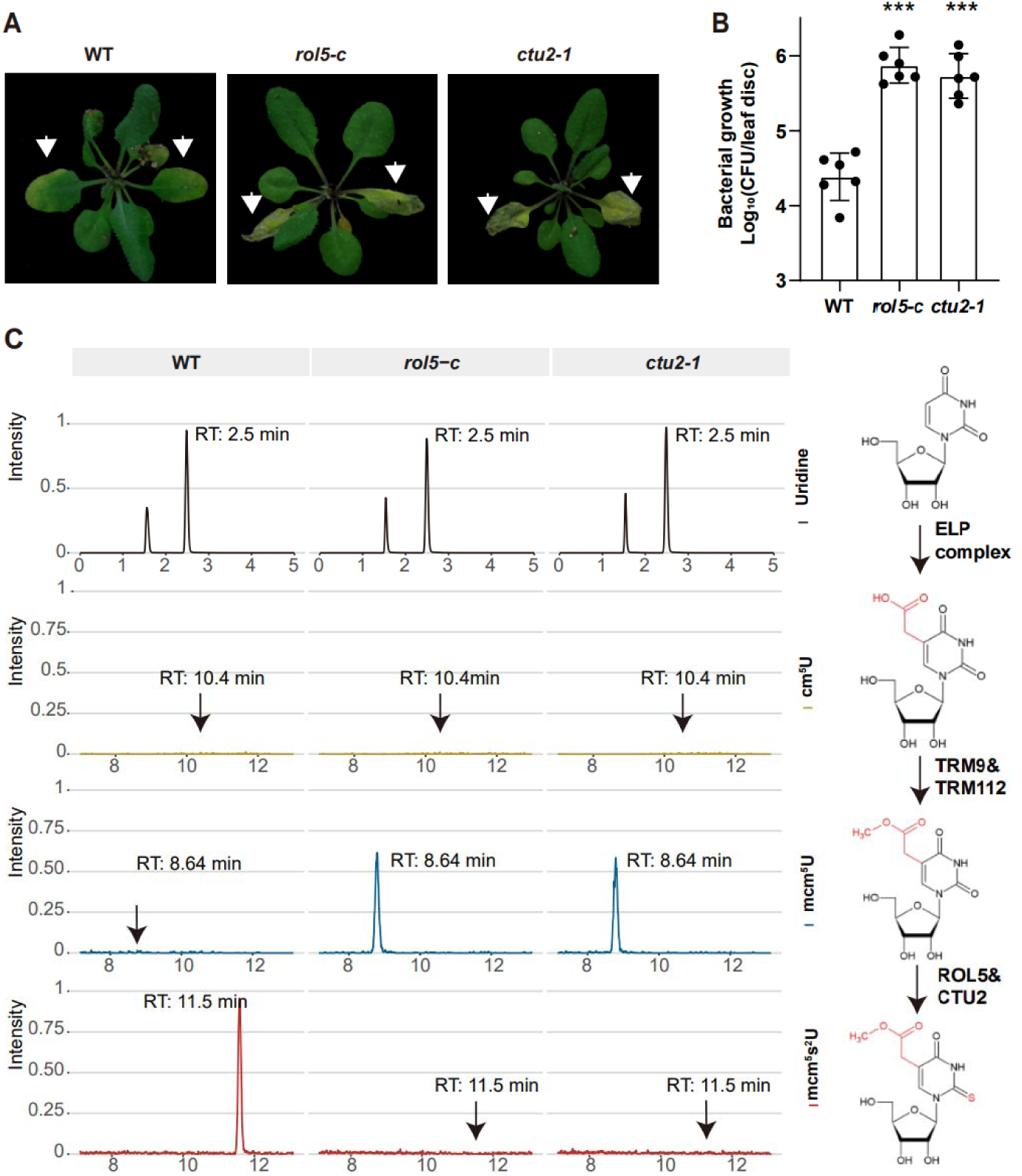
ROL5 and CTU2 is required for mcm^5^s^2^U and plant immunity. (A and B) The *rol5-c* and *ctu2-1* mutants are more susceptible to the bacterial pathogen *Psm* ES4326 than WT. (A) The photo of *Arabidopsis* 3 days after infection. The arrows indicate the leaves inoculated with *Psm* ES4326. (B) The growth of *Psm* ES4326. CFU, colony-forming unit. Error bars represent 95% confidence intervals (n = 6). Statistical significance was determined by two-tailed Student’s t-test. ***, p < 0.001. (C) The *rol5-c* and *ctu2-1* mutants lack the mcm^5^s^2^U modification. The levels of U, cm^5^U, mcm^5^U, and mcm^5^s^2^U were quantified through HPLC-MS analysis. The intensity and the retention time of each nucleotide are shown. The structure of each nucleotide and the catalyzing enzymes are shown on the right.

To investigate whether ROL5 and CTU2 are indeed required for mcm^5^s^2^U in plants, we measured the mcm^5^s^2^U levels in WT, *rol5-c*, and *ctu2-1* using high-performance liquid chromatography with mass spectrometry (HPLC-MS). In WT, mcm^5^U was almost undetectable (Figure 3C), indicating that mcm^5^U is efficiently transformed into mcm^5^s^2^U in *Arabidopsis*. However, in the *rol5-c* and *ctu2-1* mutants, the mcm^5^s^2^U level was undetectable while the mcm^5^U level was very high, suggesting that both ROL5 and CTU2 are required for mcm^5^s^2^U. These data revealed that ROL5 and CTU2 form a complex to catalyze the mcm^5^s^2^U modification, which is essential for plant immunity.

### The transcriptional and translational reprogramming is compromised in *cgb*

To understand why the *cgb* mutant was hyper-susceptible to pathogens, we performed transcriptome and proteome analysis of the *cgb* mutant and the *COM* line. Each sample was divided into two parts, one for transcriptome analysis using RNA sequencing (RNA-seq) approach, and the other for proteome analysis using tandem mass tag (TMT)-based approach. Principal Component Analysis (PCA) showed that the reproducibility between biological replicates was good (Supplemental Figure S1). The differentially expressed genes (DEGs) and the differentially expressed proteins (DEPs) between different samples were obtained through data analysis. In *COM*, 22% (4819) and 27% (5767) of genes were respectively up-regulated or down-regulated after *Psm* infection (Figure 4A; Supplemental Table S1). However, only 18% (3986) and 23% (4913) of genes were respectively up-regulated or down-regulated in *cgb* (Supplemental Table S1). In *COM*, 16% (1193) and 13% (1021) of proteins were respectively up-regulated or down-regulated after *Psm* infection (Figure 4B; Supplemental Table S2). In contrast, only 12% (909) and 10% (787) of proteins were respectively up-regulated or down-regulated in *cgb* (Supplemental Table S2). Therefore, the numbers of both DEGs and DEPs in *cgb* were less than those in *COM*.

**Figure 4.**
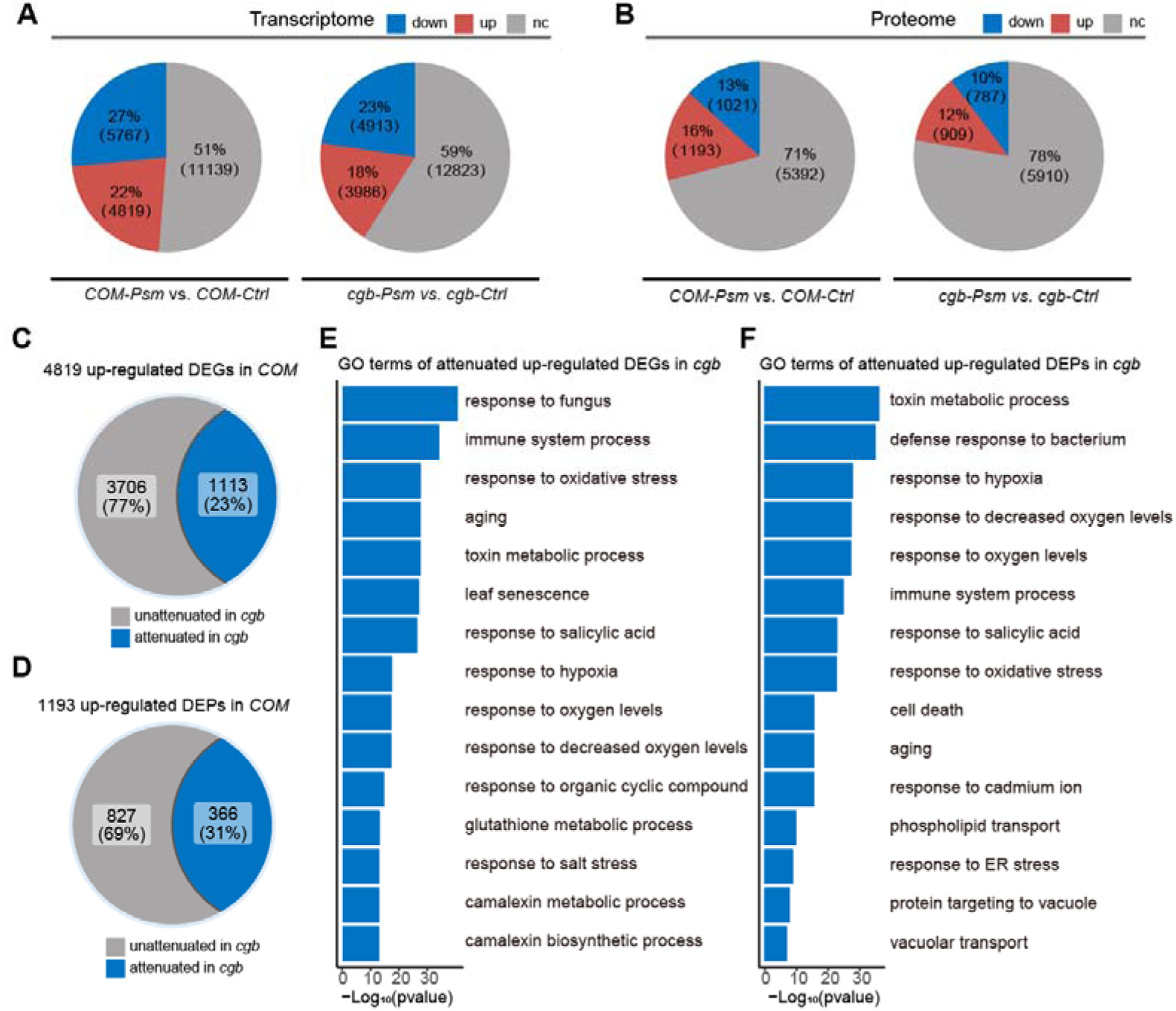
The transcriptional and translational reprogramming is compromised in *cgb.* (A and B) The percentage and the number of the differential expressed genes (DEGs, *p*-value < 0.05, |Log_2_Foldchange| > Log_2_1.5, A) and the differential expressed proteins (DEPs, *p*-value < 0.05, |Log_2_Foldchange| > Log_2_1.2, B) after *Psm* infection in the *cgb* mutant and the complementation line (*COM*). Down, down-regulated. Up, up-regulated. Nc, no change. (C and D) The percentage and the number of the attenuated genes (C) and proteins (D) in *cgb* among the up-regulated DEGs and DEPs in *COM*. (E and F) Gene Ontology (GO) analysis of the attenuated genes or proteins in *cgb*. The top 15 significantly enriched GO terms are shown.

To further examine the gene expression defects in *cgb*, we compared the expression changes after *Psm* infection between *cgb* and *COM*. Among 4819 up-regulated DEGs in *COM*, the expression changes of 1113 genes were less prominent in *cgb* than in *COM* (Figure 4C; Supplemental Table S3). These genes were referred to as attenuated genes. Among 1193 up-regulated DEPs in *COM*, the expression changes of 366 proteins were less prominent in *cgb* than in *COM* (Figure 4D; Supplemental Table S4). These proteins were named attenuated proteins. Gene Ontology (GO) analysis of the attenuated genes and attenuated proteins revealed that many important biological processes were significantly enriched (Figure 4E and F; Supplemental Table S5). These data suggested that both transcriptional reprogramming and translational reprogramming were compromised in *cgb*.

### The translation efficiency of immune-related proteins is compromised in *cgb*

Since the mcm^5^s^2^U modification directly regulates translation process, we sought to identify the proteins with compromised translation efficiency. The 366 attenuated proteins in *cgb* may be due to reduced transcription or reduced translation. To distinguish these two possibilities, we performed venn diagram analysis between attenuated genes and attenuated proteins, revealing that 261 attenuated proteins were not attenuated genes, suggesting that the attenuated expression of these proteins is due to reduced translation (Figure 5A; Supplemental Table S6). GO analysis of these 261 proteins revealed that the immune-related processes (i.e., Response to Salicylic Acid, Defense Response to Bacterium, and Immune System Process) were significantly enriched (Figure 5B; Supplemental Table S7). Notably, NPR1 is one of these proteins.

**Figure 5.**
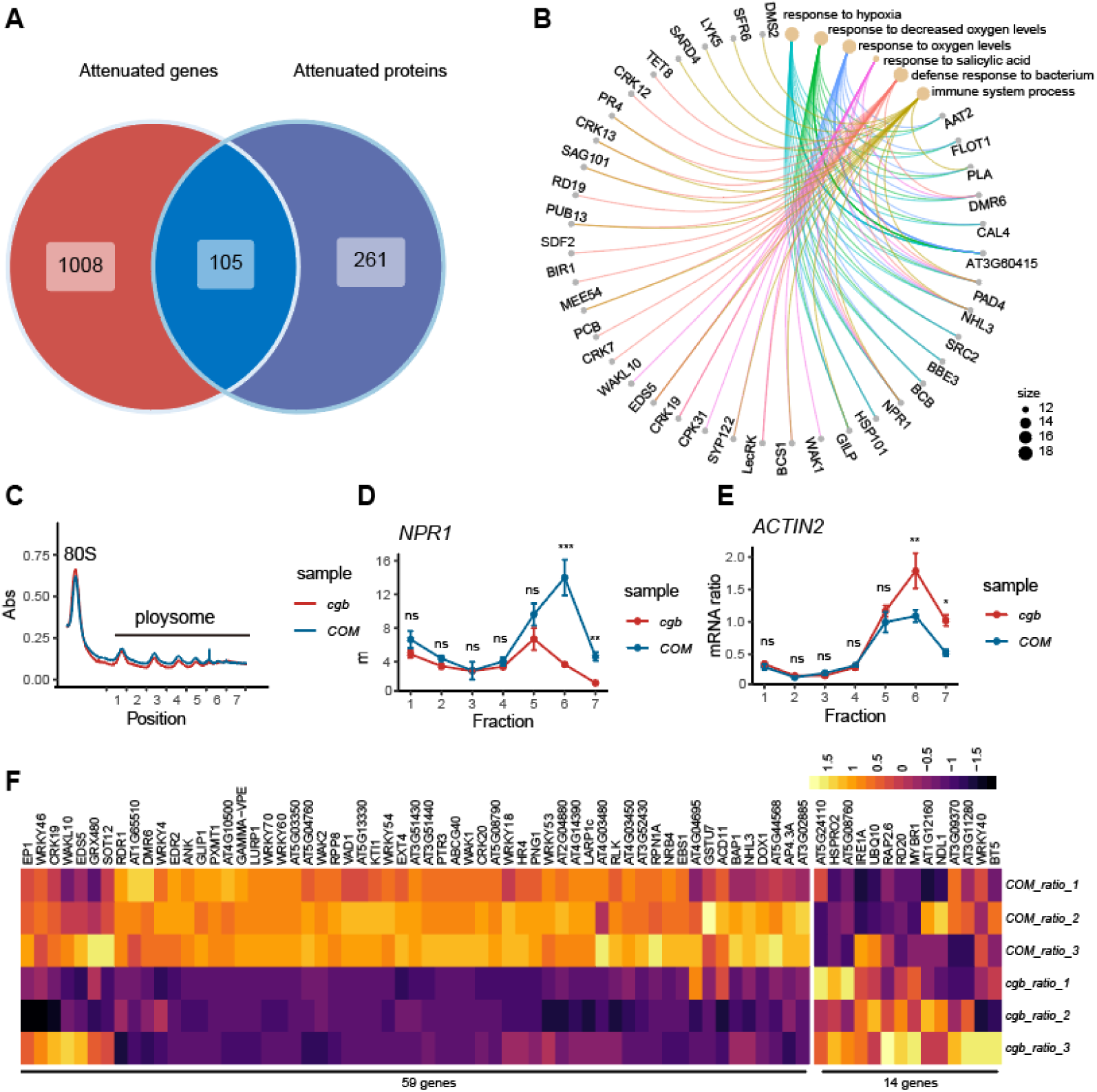
The translation of immune-related proteins is compromised in *cgb.* (A) Venn diagram analysis of attenuated genes and proteins. (B) GO analysis of the 261 attenuated proteins. The top 6 significantly enriched GO terms are shown. (C) Polysome profiling results. Abs, the absorbance of sucrose gradient at 254 nm. The numbers on X-axis indicate the polysomal fractions subjected for the qPCR analysis. (D and E) The qPCR analysis results. The relative mRNA level of each gene in different fractions or in total mRNA were normalized against UBQ5. The ratio between the relative mRNA levels in each fraction and in total mRNA were shown (n = 3). Statistical significance was determined by two-tailed Student’s t-test. *, p < 0.05; **, p < 0.01; ***, p < 0.001; ****, p < 0.0001; ns, no significance. (F) The heatmap showing the expression changes of SA-responsive genes after pathogen infection.

To confirm that the translation of NPR1 was reduced in *cgb*, we carried out ribosome profiling experiment. Compared with *COM,* the polysome fractions in *cgb* were reduced (Figure 5C), suggesting that the overall translation efficiency is lower in *cgb*. As expected, the relative mRNA levels of NPR1 in multiple polysome fractions were significantly lower in *cgb* than in *COM* (Figure 5D). It is of note that the negative control ACTIN2 was not reduced but was enhanced in *cgb* (Figure 5E).

The reduced NPR1 protein level in *cgb* suggested that SA signaling is compromised. To test this possibility, we examined the expression of all genes (118) belonging to the GO term Response to Salicylic Acid. In our transcriptome data, we could detect the expression of 73 genes, among which 59 genes (80.8%) were reduced in *cgb* compared with *COM* (Figure 5F; Supplemental Table S8). Therefore, SA signaling is indeed comprised in *cgb*, which may be one of the major reasons why *cgb* was hypersusceptible to pathogen.

## Discussion

Upon pathogen infections, plants need to efficiently reprogram their gene expression, allowing the transition from growth to defense. However, how the translation system contributes to the immune response is not well-studied. The tRNA thiolation is required for efficient protein expression (Schaffrath and Leidel, 2017; Nedialkova and Leidel, 2015). In the *cgb* mutant, the tRNA thiolation was abolished (Figure 3), leading to disease hyper-susceptibility (Figure 1). We found that the translation of many immune-related proteins was reduced in *cgb* (Figure 5; Supplemental Table S6). Therefore, our study strongly suggested that tRNA thiolation is required for plant immunity, thus revealing a new mechanism underlying plant immune responses. Nevertheless, how plants and pathogens regulate tRNA thiolation remains to be further studied.

The SA receptor NPR1 is the master regulator of SA signaling. NPR1 can function as a transcription coactivator to regulate gene expression or an adaptor of ubiquitin E3 ligase to mediate protein degradation (Yu et al., 2021; Zavaliev et al., 2020; Yu et al., 2022). It has been shown that the activity of NPR1 is regulated at multiple levels including post-translational modifications such as phosphorylation, ubiquitination, S-nitrosylation, and sumoylation (Saleh et al., 2015; Spoel et al., 2009; Tada et al., 2008). However, how NPR1 is regulated at the translational level is unknown. Here we show that the tRNA thiolation-mediated translation control is required for the optimal expression of NPR1 (Figure 5B and 5D), revealing a new layer of regulation for NPR1. It will be interesting to study further how tRNA thiolation regulates NPR1 translation.

The tRNA thiolation is highly conserved in eukaryotes. However, its biological functions in plants are less well-understood. Previously, it was reported that tRNA thiolation regulates the development of root hair, chloroplast, and leaf cells (Philipp et al., 2014; Leiber et al., 2010). Recently, it was found that tRNA thiolation is required for heat stress tolerance (Xu et al., 2020). Our study found that tRNA thiolation is involved in plant immunity, thereby revealing a new biological function of tRNA thiolation. It will be interesting to test whether tRNA thiolation is required for responses to other stresses such as drought, salinity, and cold.

It is generally believed that the stress-responsive genes play important roles in stress responses. And many studies have identified numerous pathogen-responsive genes through transcriptome analysis (Zhang et al., 2020). However, previous studies have shown that the correlation between mRNAs and proteins is not strong (Schwanhüusser et al., 2011; Lahtvee et al., 2017). Given that proteins are major players in cellular functions, it is necessary to study immune responses at the protein level. Through high-throughput proteome analysis, we found that 2215 proteins are differentially expressed after *Psm* infection in *Arabidopsis* (Figure 4A; Supplemental Table S2). To our knowledge, this is the largest dataset of pathogen-responsive proteins in *Arabidopsis*. We believe that this dataset will be a good research resource for future studies on plant immunity.

## Materials and Methods

### Plant material and growth conditions

All *Arabidopsis* seeds used in this study are in *Columbia-0* background. The *npr1-1* mutant was described previously (Cao et al., 1997). The *cgb* mutant and the complementation line were generated in this study. The mutant of *ctu2-1* (SALK_032692) was purchased from ABRC. The *rol5-c* mutant was generated using EC1-based CRISPR-Cas9 system (Wang et al., 2015). All seeds were sterilized with 2% Plant Preservative Mixture-100 (Plant Cell Technology) at 4°C in the dark for 2 days and then were plated on Murashige and Skoog (MS) medium with 1% sucrose and 0.3% phytagel. The plants were grown under long-day conditions at 22°C (16 h of light/8 h of dark; supplied by white-light tubes).

### Strains and growth conditions

*E. coli* strain *DH5*α for molecular cloning was cultured in LB medium at 37°C. *E. coli* strain *BL21* (DE3) for recombinant protein expression was cultured in LB medium at 16°C. *Agrobacterium tumefaciens* strain GV3101 for transformation was cultured in Yeast Extract Beef (YEB) medium at 28°C. *Psm* ES4326 and *Pst* DC3000 avrRPT2 for infection assay was cultured in King’s B (KB) medium at 28°C. Yeast strain AH109 for yeast-two-hybrid assay was cultured in Yeast Peptone Dextrose (YPD) medium or SD medium at 28°C.

### Vector constructions

The vectors were constructed using digestion–ligation method or a lighting cloning system (BDIT0014, Biodragon Immunotechnology). For complementation experiment, *ROL5* was inserted into *Nco* I/*Xba* I-digested *pFGC5941*. For pull-down assays, *CTU2* was inserted into *Bam*H I/*Xho* I-digested *pGEX-6P-1*; *ROL5* was inserted into *Nco* I/*Hin*d III-digested pET28a. For split luciferase assays, *ROL5* and *CTU2* were cloned into the *Kpn* I/*Sal* I-digested *pJW771* and *pJW772*, respectively. For yeast two hybrids assays, *ROL5* and *CTU2* were cloned into *EcoR* I/*BamH* I-digested *pGBKT7* and *pGADT7*, respectively. For the co-immunoprecipitation assay, *ROL5-FLAG* and *CTU2-GFP* were cloned into *Nco* I/*Xba* I-digested *pFGC5941*. To generate *rol5-c*, the target sequence was designed and cloned into *pHEE401* as described previously(Wang et al., 2015). The primer sequences used for cloning were listed in Supplemental Table S9.

### Reverse transcription and PCR

The total RNA or the RNA in ribosome fractions were extracted using was extracted using TRIzol Reagent (Invitrogen). The cDNA was synthesized using HiScript II Q RT SuperMix (Vazyme, China). The qPCR analysis was performed using the AceQ qPCR SYBR Green Master Mix (Vazyme Biotech, Nanjing, China). UBQ5 were used as the internal reference gene. Primers used for qPCR were listed in Supplemental Table S9.

### Pathogen infection

The third and fourth leaves of the three-week-old *Arabidopsis* were infiltrated with *Psm* ES4326 (OD_600_ = 0.0002) or *Pst* DC3000 avrRPT2 (OD_600_ = 0.02) using a needleless syringe.

### Yeast-two-hybrid assays

The bait and prey vectors were co-transformed into the yeast strain AH109. Protein-protein interactions were determined by yeast growth on SD/-Leu/-Trp/-His/ medium.

### *In vitro* pull-down assays

The GST pull-down assays were performed as previously described (Pan et al., 2021). Briefly, the ROL5-His, GST, and GST-CTU2 proteins were expressed in *E. coli* BL21 (DE3). GST and GST-CTU2 were coupled to Glutathione beads (GE Healthcare Life Sciences) and then were incubated with ROL5-His in the binding buffer (50 mM Tris–HCl pH 7.5, 150 mM NaCl, 1 mM EDTA, and 2 mM DTT) at 4°C for 2 h. The beads were washed three times with washing buffer (binding buffer plus 2% NP-40), boiled in 1× SDS loading buffer, and analyzed by western blot using anti-GST antibody (ABclonal).

### Co-immunoprecipitation assays

The co-immunoprecipitation assays were performed as previously described(Chen et al., 2021). 35S:*ROL5-FLAG* and 35S:*CTU2-GFP* were transformed into *Agrobacterium tumefaciens* GV3101. 35S*:ROL5-FLAG* strain was infiltrated alone or co-infiltrated with 35S*:CTU2-GFP* into the leaves of *N. benthamiana*. After 48 h, the infiltrated leaves were ground in liquid nitrogen and were resuspended in IP buffer (20 mM Tris–HCl pH 7.5, 50 mM NaCl, 1 mM EDTA, 0.1% SDS, 1% Triton X-100, 1 mM PMSF, 100 μM MG132, 1× protease inhibitor cocktail) for total protein extraction. The lysates were incubated with GFP-Trap magnetic beads (Chromotek) at 4°C for 2 h. The beads were washed using washing buffer (20 mM Tris–HCl pH 7.5, 150–500 mM NaCl, 1 mM EDTA, 1 mM PMSF, 1× Protease Inhibitor Cocktail) and then boiled in 1× SDS loading buffer. The western blotting was performed using anti-FLAG (Promoter) and anti-GFP (Promoter) antibodies.

### Split luciferase assays

Split luciferase assay was performed as described previously(Chen et al., 2008). The constructs were transformed into *Agrobacterium tumefaciens* strain GV3101. The resultant strains were then infiltrated into leaves of *N. benthamiana.* After 48 h, 1 mM luciferin (GOLDBIO) was applied onto leaves and the images were captured using Lumazone imaging system equipped with 2048B CCD camera (Roper).

### Quantification of tRNA modifications

Quantification of tRNA modifications was performed using liquid chromatography coupled with mass spectrometry (LC/MS) according to a previous study(Su et al., 2014). Total tRNA was extracted using a microRNA kit (Omega Bio-Tek, Norcross, USA). Five micrograms of tRNA were hydrolyzed in 10 μL enzymic buffer (1 U Benzonase, 0.02 U Phosphodiesterase I, and 0.02 U Alkaline phosphatase) at 37°C for 3 h. The UHPLC system (Thermo Fisher Scientific, Waltham, USA) coupled with TSQ Altis Triple Quadrupole Mass Spectrometer (Thermo Fisher Scientific, Waltham, USA) was used for quantification of tRNA modification. For the liquid chromatography, the Hypersil GOLD^TM^ HPLC column (3 µm, 150 × 2.1 mm; Thermo Fisher Scientific) was used. The solvent gradient was set as the protocol (Su et al., 2014). The Tracefinder software (Thermo Fisher Scientific, Waltham, USA) was further used for peak assignment, area calculation, and normalization. Corresponding structures and molecular masses were obtained from the Modomics database (https://iimcb.genesilico.pl/modomics/modifications).

### RNA and protein extraction for transcriptome and proteome analysis

The samples were grinded in liquid nitrogen and divided into two parts, one for transcriptome analysis and the other for proteome analysis. Total RNA was extracted using Trizol reagent (Aidlab, Beijing, China). Library preparation and RNA-sequencing were performed by Novogene (Beijing, China). Total proteins were extracted using phenol-methanol method (Deng et al., 2007). The protein concentration was determined with 2-D Quant Kit (GE Healthcare Life Sciences, Westborough, USA) using Bovine Serum Albumin (BSA) as a standard.

### Proteome analysis

For trypsin digestion, 60 μg proteins of each sample were reduced with 20 mM Tris-phosphine for 60 min at 30°C. Cysteines were alkylated with 30 mM iodoacetamide for 30 min at room temperature in the dark. Proteins were precipitated with 6 volumes of cold acetone overnight and then dissolved in 50 mM TEAB. Proteins were digested with trypsin (protease/protein = 1/25, w/w) overnight at 37°C.

For TMT labeling, each sample containing 25 μg of peptide in 50 mM TEAB buffer was combined with its respective 10-plex TMT reagent (Thermo Fisher Scientific) and incubated for 1 h at room temperature. Three biological replicates were labeled respectively for each sample, in which the samples of *COM* were labeled with 126-, 127N- and 128C- of the 10-plex TMT reagent, while the samples of *cgb* were labeled with 129N-, 130C-, and 131- of the 10-plex TMT reagents. The labeling reactions were stopped by the addition of 2 μL of 5% hydroxylamine.

For LC-MS/MS analysis, multiplexed TMT-labeled samples were combined, vacuum dried, and reconstituted in 2% acetonitrile and 5 mM ammonium hydroxide (pH 9.5), and separated with the Waters Acquity BEH column (C18, 1.7μm, 100mm, Waters) using UPLC system (Waters) at a flow rate 300 μl/min. Total of 24 fractions were collected, combined into 12 fractions, and vacuum dried for LC-MS/MS analysis. The solvent gradient was set as previously described (Deng et al., 2007). Samples were then analyzed on an Ultimate 3000 nano UHPLC system (Thermo Fisher Scientific) coupled online to a Q Exactive HF mass spectrometer (Thermo Fisher Scientific). The trapping column (PepMap C18, 100Å, 100 μm×2 cm, 5μm) and an analytical column (PepMap C18, 100Å, 75 μm i.d.×50 cm long, 2μm) were used for separation of the sample. The solvent gradient and MASS parameters were set as previously described (Deng et al., 2007).

### Transcriptome data analysis

Raw reads were processed and aligned to the *Arabidopsis* genome (https://www.arabidopsis.org) using STAR (v.2.6.1a). Genes with over 10 reads were filtered and processed using DESeq2 (v.1.22.2) to identify the differentially expressed genes (*p*-value < 0.05, |Log_2_FoldChange| > Log_2_1.5) (Love et al., 2014).

### Proteome data analysis

Raw data were processed using Proteome Discoverer (v.2.2.0.388) and aligned to *Arabidopsis* genome (https://www.arabidopsis.org) with the SEQUEST HT search engine. Searches were configured with static modifications for the TMT reagents (+229.163 Da). The precursor mass tolerance was set as 10 ppm; the fragment mass tolerance was set as 0.02 Da; the trypsin missed cleavage was set as 2. The reversed sequence decoy strategy was used to control peptide false discovery. The peptides with q scores < 0.01 were accepted, and at least one unique peptide was required for matching a protein entry for its identification. PSMs (peptide spectrum matches) results were processed with DESeq2 (v.1.22.2) to identify the differentially expressed proteins (*p*-value < 0.05, |Log_2_FoldChange| > Log_2_1.2).

### GO and Heatmap analysis

The differentially expressed genes or proteins were analyzed by using Clusterprofile (v.3.18.1) (Yu et al., 2012). The heatmap analysis was processed by using pheatmap2 (v.1.0.12).

### Ribosome profiling

The ribosome profiling was performed as the previous description with some modifications (Hsu et al., 2016; Xu et al., 2017). The plant sample (0.05-0.1 g) was grinded in liquid nitrogen and extracted with 1 mL ribosome lysis buffer (200 mM Tris-HCl pH 8.0, 200 mM KCl, 35 mM MgCl_2_, 1% Triton X-100, 100 μM MG132, 1 mM DTT, and 100 μg/mL cycloheximide), followed by ultracentrifugation at 4°C for 2 h (38000 rpm, Beckman, SW41 rotor) through a 20-60% sucrose gradient (40 mM Tris-HCl pH 8.4, 20 mM KCl, 10 mM MgCl_2_, and 50 μg/mL cycloheximide) prepared by Gradient Master (Biocomp Instruments, Fredericton, Canada). The profiling signals were recorded by Piston Gradient Fractionator (Biocomp Instruments, Fredericton, Canada).

### SA quantification

SA extraction and LC/MS manipulation were performed as previously described with some modifications (Kusch et al., 2019). Plant material (0.1 g) was grinded in liquid nitrogen and were extracted by using 3 mL Methyl-tert-butyl ether/methanol/water (10:3:2.5, V/V) containing 10 ng D4-SA (CDN Isotopes, Pointe-Claire, Canada) for 1 h at room temperature. The upper phase was then collected and dried by the vacuum dryer for the LC/MS analysis. The UHPLC coupled with TSQ Altis Triple Quadrupole Mass Spectrometer was used to detect the relative concentration of SA (Thermo Fisher, Waltham, USA). Solvent A and B (pH 3.5) were water and acetonitrile/water (90:10, v/v), respectively. The flow rate was 0.16 mL min−1 and the separation temperature was constantly at 40 °C. Column Hyersil GLOD (Thermo Scientific, cat. no.25005-152130) was used to separate with the UHPLC system. The LC/MS method was performed as previously described (Kusch et al., 2019). The Tracefinder software (Thermo Fisher, Waltham, USA) was used for peak assignment, area calculation, and normalization.

### Ion leakage assay

The third and fourth leaves of the three-week-old Arabidopsis were infiltrated with *Pst* DC3000 carrying avrRPT2 (OD_600_ = 0.02) using a needleless syringe. For measurements of ion leakage, 8 leaf discs were floated in 50 mL of ultrapure water. The conductivity was measured every 4 h using a conductivity meter (FiveGo^TM^ F3, Mettler Toledo, Columbus, USA).

### Data availability

RNA sequencing datasets have been deposited to GSE database (https://www.ncbi.nlm.nih.gov/geo/) with an accession number GSE183087. The mass spectrometry proteomics data have been deposited to the ProteomeXchange Consortium (http://proteomecentral.proteomexchange.org) via the iProX partner repository with the dataset identifier PXD028189. Data analysis scripts are available on GitHub: https://github.com/XueaoZHENG/cgb_project.

## Accession numbers

Sequence data from this article can be found in the *Arabidopsis* Genome Initiative or GenBank/EMBL databases under the following accession numbers: ROL5(AT2G44270), CTU2(AT4G35910).

## Supplemental data

**Supplemental Figure S1.** Principal Component Analysis (PCA) of the transcriptome and proteome samples.

**Supplemental Table S1.** The differentially expressed genes after *Psm* infection in *cgb* mutant and its complementation line.

**Supplemental Table S2.** The differentially expressed proteins after *Psm* infection in *cgb* mutant and its complementation line.

**Supplemental Table S3.** The attenuated genes in *cgb*.

**Supplemental Table S4.** The attenuated proteins in *cgb*.

**Supplemental Table S5.** The enriched GO terms of attenuated genes and proteins.

**Supplemental Table S6**. The attenuated proteins with reduced translation in *cgb*.

**Supplemental Table S7**. The enriched GO terms of the attenuated proteins with reduced.

**Supplemental Table S8**. The gene expression changes of SA-responsive genes after pathogen infection.

**Supplemental Table S9**. The primers used in this study.

## Acknowledgements

We are grateful to Dr. Zhipeng Zhou and Dr. Peng Chen for helpful suggestions and discussion. We thank BaiChuan Program from College of Life Science and Technology, Huazhong Agricultural University.

## Author contributions

S.Y., and X.Z. designed the project. H.C. and C.W. identified ROL5. H.C. found the interaction between ROL5 and CTU2. X.Z. and L.Z. performed quantification of mcm^5^s^2^U. H.C. and Y.W. performed pathogen infection assay. X.Z. performed RNA-seq assay. Z.D. and X.Z. performed TMT-based proteome assay. X.Z. performed bioinformatic analysis. X.Z. and S.Y. wrote the manuscript with inputs from others.

