## Supplemental Figure S1 for "The tRNA thiolation-mediated translational control is essential for plant immunity"

### Supplementary Figures

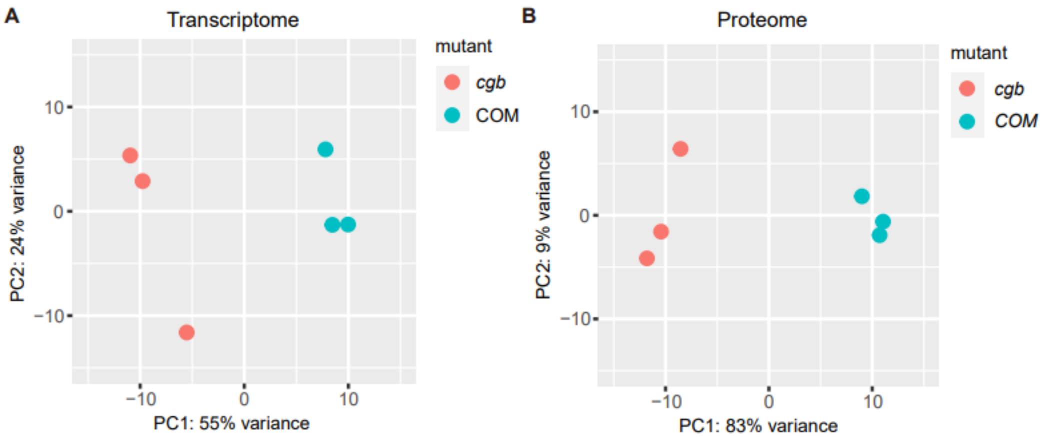


**Figure S1.** Principal Component Analysis (PCA) of the transcriptome and proteome samples
